# De novo design of functionally diverse and druggable antimicrobial peptides by diffusion and multimodal deep learning

**DOI:** 10.1101/2025.07.16.665042

**Authors:** AnQi Ouyang, Fang Ge, Jiangning Song, Dong-Jun Yu

## Abstract

The escalating crisis of antibiotic resistance underscores the urgent need for innovative anti-infective agents, such as antimicrobial peptides (AMPs), though their discovery and optimization remain challenging. To address this, we introduce AMP-D3, a comprehensive framework that integrates AMP generation, identification, and screening into a cohesive workflow. Central to AMP-D3 is PACD, an advanced generator that utilizes contrastive diffusion to produce diverse, functionally tailored AMPs. For precise recognition, the CAST model combines ESM-2 embeddings with localized sequence features through cross-modal attention, yielding an accuracy of 0.891—a 2.2% enhancement over current state-of-the-art approaches. Additionally, a robust multi-attribute prediction module evaluates key properties, including antifungal and anticancer potential, across twenty-two distinct characteristics. Furthermore, a meticulously designed screening pipeline identifies high-efficacy AMP candidates against critical pathogens, such as *C. albicans, E. coli, P. aeruginosa*, and *S. aureus*. This process assessed 10,000 peptides generated by PACD alongside 43,000 from prior studies, selecting the top 100 candidates per pathogen for further analysis. Experimental validation confirms that these peptides exhibit structural diversity and binding affinities comparable to naturally occurring AMPs, as evidenced by AlphaFold2 structural predictions and molecular docking studies. Thus, AMP-D3 represents a transformative approach to the design and expedited discovery of clinically viable AMPs.

## Introduction

In recent years, the surge in antibiotic-resistant bacterial strains due to antibiotic misuse has posed a severe challenge to global public health security. Resistant infections not only impose a significant clinical burden but also exert substantial socioeconomic pressure, making the development of novel anti-infective drugs a priority in biomedical research ^1^. Antimicrobial peptides (AMPs), a class of therapeutic polypeptides composed of short amino acid chains, have demonstrated significant potential in treating infectious diseases due to their unique bactericidal mechanisms and low propensity for inducing resistance^2^. Unlike traditional antibiotics, AMPs offer multiple advantages: they target bacterial cell membranes, forming transmembrane ion channels that disrupt membrane integrity and cause leakage of cellular contents, thereby exerting their bactericidal effects ^3^. This non-specific mechanism makes it difficult for bacteria to develop resistance through missense mutations, significantly reducing the risk of resistance ^4^.

Traditionally, AMPs discovery has relied on two main strategies: (1) experimental screening based on phage/yeast display technologies ^5^, and (2) systematic evaluation of sequence space using computational tools to identify candidate peptides with specific functional properties^6^. However, the vast combinatorial space of peptide sequences, coupled with the rarity of clinically valuable active peptides, renders traditional screening methods inefficient and costly. Moreover, AMPs face several challenges in clinical applications, including high cytotoxicity ^7^, sensitivity to extreme conditions (e.g., pH and temperature) ^8^, insufficient target specificity, folding stability issues in large AMPs ^9^, microbial resistance ^10^, and high production costs ^11^. Recent breakthroughs in deep learning for protein structure prediction ^12^ have opened new avenues for AMPs research, significantly advancing progress in rational design, efficient identification, activity prediction, and performance optimization.

Three transformative developments characterize contemporary AMP research: First, deep learning has supplanted conventional machine learning in identification tasks. Early predictors like PPTPP ^13^and Macrel ^14^ utilized random forests with handcrafted features, whereas modern architectures employ adaptive feature learning through CNNs (iAMPCN ^15^), RNNs (AMPlify ^16^), and Transformer (AMP-BERT ^17^). Notably, Diff-AMP ^18^ leverages the ESM-2 ^19^ to capture long-range amino acid dependencies, demonstrating enhanced generalization through transfer learning. Second, deep generative models have enabled de novo AMP design by learning latent peptide distributions - approaches range from GAN variants (AMPGANv2 ^20^) and VAE frameworks ^21^ to emerging diffusion models ^22^ with superior distribution-fitting capacities ^23^. Third, multi-task learning architectures now predict diverse bioactivities, exemplified by TransImbAMP_’_s seven-label classification ^24^ and iAMP-CA2L_’_s cellular automata-based prediction of ten activities ^25^.

Current AMP generation technologies encounter four key limitations: (1) Incomplete use of therapeutic and non-therapeutic peptide data, reducing model accuracy. (2) Inefficient feature extraction in screening processes, causing elevated false-negative rates. (3) Lack of attribute-driven generation for specific clinical applications, such as antiviral or anticancer therapies. (4) Challenges in assessing critical pharmacological traits—stability, toxicity, and hemolytic activity—vital for clinical success. These shortcomings obstruct precision medicine goals in AMP research.

To address these critical limitations, we propose AMP-D3, a cohesive framework that merges contrastive diffusion and multimodal fusion to streamline AMP development by enabling de novo peptide generation, precise AMP identification, broad-spectrum activity prediction, and comprehensive druggability screening. Our primary contributions are as follows: (1) We developed PACD: for *de novo* AMP generation. This cutting-edge AMP generator that leverages contrastive diffusion principles to amplify sequence diversity while preserving therapeutic efficacy. (2) We designed CAST: for precise AMP identification. This advanced multi-head attention architecture that fuses raw sequence data with high-dimensional semantic embeddings, thereby enhancing screening precision. (3) We constructed a multi-attribute prediction module: for comprehensive broad-spectrum activity prediction. This tailored component designed to adapt AMP functionality to specific clinical needs. (4) We established a targeted antimicrobial peptide screening pipeline: for efficient druggable peptide screening. This systematically structured pipeline crafted to pinpoint highly potent AMPs against key pathogens, including *C. albicans, E. coli, P. aeruginosa*, and *S. aureus*. This pipeline incorporates an antimicrobial activity ranking module and a druggability metric screening module, ensuring rigorous and accurate selection. This integrated pipeline redefines intelligent AMP design by seamlessly connecting computational pattern recognition with practical therapeutic outcomes.

## Results

### AMP-D3 Architecture

We developed AMP-D3, an integrated framework synergistically combining deep generative modeling, hybrid feature recognition, and multi-task prediction for accelerated antimicrobial peptide development. As illustrated in **Fig. 1**, this architecture operates through four functionally chained modules, as follows: (1) PACD: the contrastive diffusion generator (**Fig. 1A**) produces evolutionarily plausible candidate AMPs (c-AMPs) via noise-perturbation denoising cycles. (2) CAST: the hybrid feature recognizer (**Fig. 1B**) implements attention-based fusion of ESM-2 embeddings ^19^ with handcrafted sequence descriptors for precise identification. (3) A multi-attribute prediction module (**Fig. 1D**) enables concurrent evaluation of 22 clinically relevant bioactivities through parallel feature extraction pathways. (4) Schematic of the screening pipeline for highly active and druggable AMPs (**Fig. 1E**). Subsequent sections provide rigorous mathematical formulations and architectural specifications for each component.

**Fig. 1:**
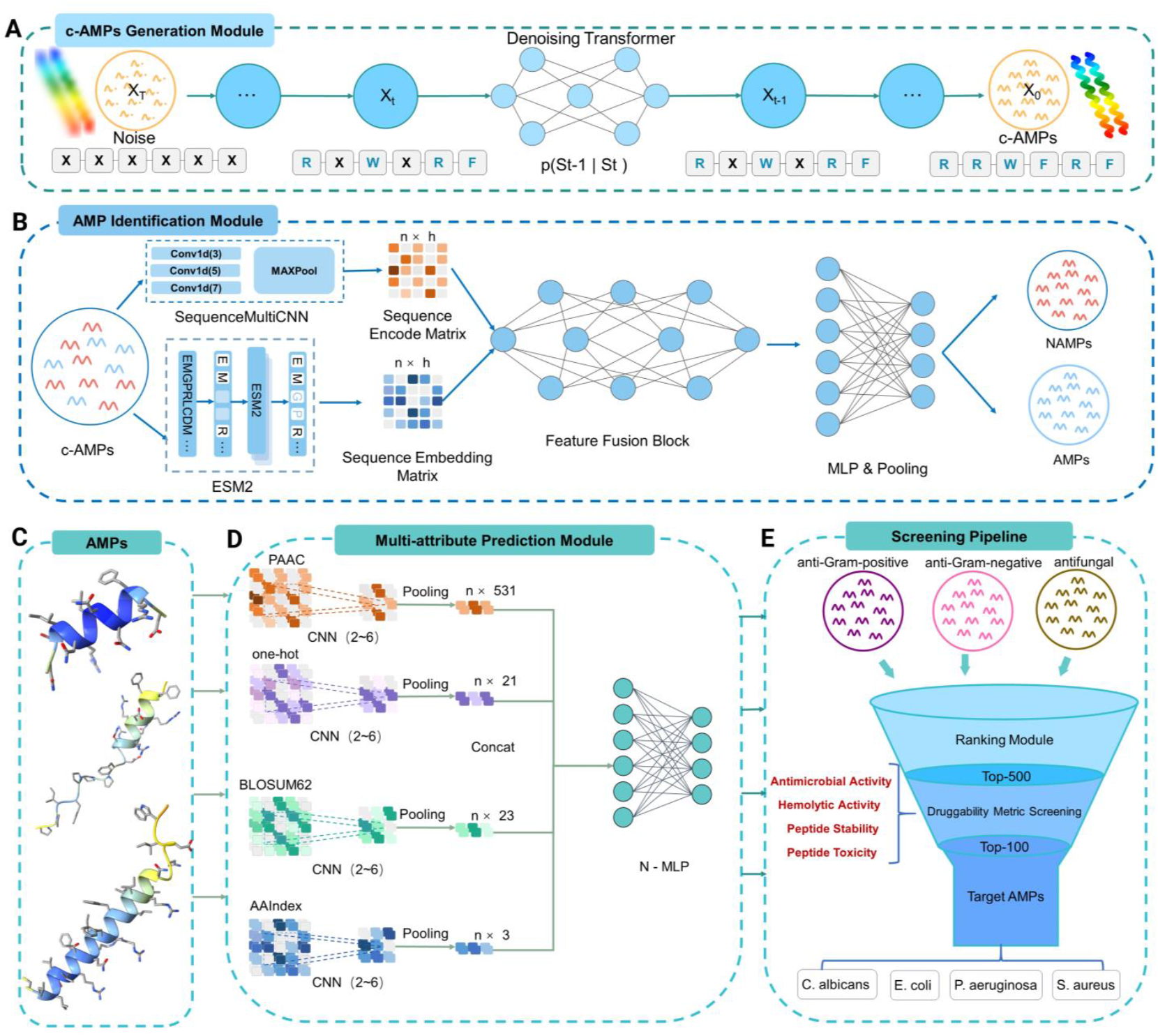
Architectural overview of AMP-D3 framework. **(A)** Contrastive diffusion generator: c-AMPs generation module based on a contrastive diffusion model, which generates high-quality candidate sequences through a noise perturbation-denoising mechanism and contrastive learning strategy. **(B)** Hybrid feature recognizer: an AMP identification module that integrates raw sequence features with deep embedding features, combining shallow sequence characteristics and deep semantic representations for precise AMPs classification. **(C)** Candidate AMP filtration: quality-controlled output from recognition module. **(D)** A multi-attribute prediction module for AMPs based on a multi-channel CNN, which comprehensively predicts multiple bioactivity metrics of AMPs through parallel feature extraction channels. **(E)** Schematic of the targeted antimicrobial peptide screening Pipeline: this pipeline processes AMPs validated by the CAST model—a sophisticated tool leveraging pattern recognition and bioinformatics—to exhibit activity against Gram-negative bacteria, Gram-positive bacteria, or fungi. These AMPs are then assessed against a panel of key pathogens, including *C. albicans, E. coli, P. aeruginosa*, and *S. aureus*. For this evaluation, a ranking module and a MIC regression model are integrated to predict antimicrobial efficacy, thereby enabling the selection of the top 100 AMPs with the highest predicted activity.

### AMPs Recognition Performance Analysis

We systematically evaluated the performance of CAST against six state-of-the-art AMP recognition approaches under identical training conditions. This benchmark included key competitors: Diff-AMP ^18^, Cao et al. ^3^, Amplify ^16^, AMPScannerV2 ^26^, MACREL ^14^, and AmPEPpy ^27^. To comprehensively assess model performance, seven standard metrics were employed: accuracy (ACC), sensitivity (SEN), specificity (SPE), precision (PRE), F1-score (F1), Matthews correlation coefficient (MCC), and area under the receiver operating characteristic Curve (AUC). Detailed descriptions and formulae for these metrics are provided in **Supplementary Note 1**.

#### Comparison with Baseline Models

CAST consistently outperformed these existing methods across all seven evaluated metrics **(Table 1)**. Notably, compared to Diff-AMP, the second-best performing model, CAST demonstrated significant improvements in ACC (+2.2%), SEN (+4.06%), MCC (+4.28%), and AUC (+0.17%). Furthermore, 10-fold cross-validation confirmed minimal data partitioning effects on CAST_’_s performance **(Table S1)**, underscoring the robustness and generalizability of the model for identifying potential antimicrobial peptides.

**Table 1.** Performance comparison of multiple AMP recognition models (%)

| Models | ACC | PRE | SEN | SPE | MCC | F1-Score | AUC |
| --- | --- | --- | --- | --- | --- | --- | --- |
| AmPEPPY | 79.99 | 87.01 | 70.50 | 89.48 | 61.09 | 77.38 | 87.41 |
| MACREL | 82.35 | 85.43 | 77.99 | 86.70 | 64.94 | 81.60 | 89.29 |
| AMPScannerV2 | 82.83 | 85.87 | 78.60 | 87.06 | 65.90 | 82.12 | 90.30 |
| Amplify | 84.16 | 88.87 | 77.89 | 90.33 | 68.84 | 83.07 | 90.69 |
| Cao et al | 85.31 | 87.92 | 81.86 | 88.75 | 70.79 | 84.71 | 91.41 |
| Diff-AMP | 86.92 | 91.48 | 83.02 | 91.44 | 74.07 | 86.39 | 92.22 |
| **Our model** | **89.12** | **92.43** | **87.08** | **91.53** | **78.35** | **89.68** | **92.39** |

CAST_’_s enhanced performance is attributed to three key design elements. First, a multi-scale feature fusion strategy integrates original peptide sequence features with high-dimensional semantic representations extracted by a pre-trained model. Second, a feature interaction module, based on multi-head cross-attention, employs a feature-wise linear modulation (FiLM) mechanism for effective features. Finally, an MLP-based feature selector automatically identifies and prioritizes key discriminative features. This end-to-end learning architecture thereby significantly enhances CAST_’_s capacity to discern and utilize crucial AMP characteristics.

#### Ablation studies for CAST

To systematically evaluate the contributions of its core components, controlled ablation experiments were conducted on the CAST architecture. These experiments compared the full CAST model against two different architectures: one lacking both the hybrid feature fusion strategy and FILM modulation (CAST w/o F-Fusion & FILM), and another without only the FILM module (CAST w/o FILM). Results indicated **(Table 2)** that the complete CAST model achieved superior performance across most evaluation metrics. The concurrent removal of feature fusion and FILM modules led to a substantial decline in performance, with ACC, SEN, MCC, and AUC decreasing by 1.4%, 5.3%, 1.8%, and 1.4% respectively, severely impairing positive sample recognition and balanced performance.

**Table 2.** Performance Comparison of Ablation Study for Antimicrobial Peptide Identification Models

| Models | ACC | PRE | SEN | SPE | MCC | F1-Score | AUC |
| --- | --- | --- | --- | --- | --- | --- | --- |
| CAST | 0.891 | 0.924 | 0.870 | 0.915 | 0.783 | 0.896 | 0.891 |
| w/o Feature-Fusion & w/o FILM | 0.877 | 0.950 | 0.817 | 0.949 | 0.765 | 0.879 | 0.877 |
| w/o FiLM | 0.886 | 0.904 | 0.884 | 0.888 | 0.771 | 0.894 | 0.886 |

The variant lacking only the FILM module also exhibited reduced efficacy, with ACC, SEN, MCC, and AUC dropping by 0.5%, 1.4%, 1.2%, and 0.5% respectively. This suggests FILM_’_s primary contribution is the enhancement of SEN, potentially by amplifying subtle antimicrobial peptide-defining patterns within fused feature representations. In conclusion, these studies demonstrate that while the feature fusion module is critical for foundational performance improvements, particularly in SEN and overall classification capability (MCC, AUC), the FILM module provides further refinement. The synergistic integration of both components is thus essential for the superior antimicrobial peptide recognition capabilities of CAST.

### AMPs Generation Quality Analysis

#### Comparison with Baseline Models

To rigorously assess generation quality, we conducted parallel experiments generating 1,000 peptide sequences using both our contrastive diffusion model and the RLGen baseline. Each model_’_s output was evaluated through five complementary metrics: ESM-2 pseudoperplexity (evolutionary plausibility), Similarity score (novelty relative to training data), Antimicrobial activity score (functional potential), BLAST identity (sequence homology), and pLDDT (structural stability). Detailed descriptions and formulae for these metrics are provided in **Supplementary Note 3**.

As demonstrated in **Table 3**, PACD significantly outperforms the RLGen model across all key metrics, highlighting its strong potential for generating therapeutic peptides. In contrast to RLGen, PACD incorporates biological knowledge into the diffusion model through contrastive learning between therapeutic and non-therapeutic peptides, thereby achieving optimal performance across various evaluation criteria. These results indicate that our model can generate more diverse and stable peptides while producing a greater number of AMPs per batch.

**Table 3.**
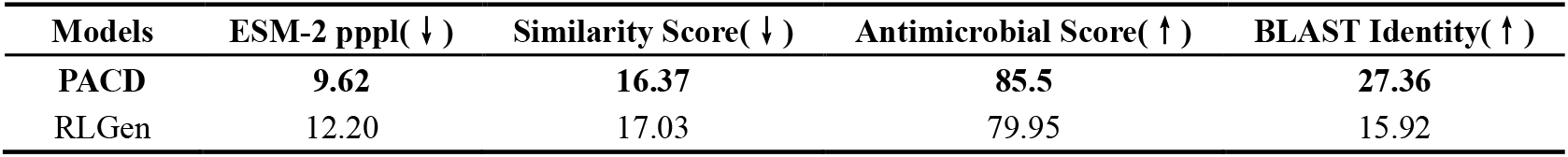
Performance comparison of PACD and RLGen architectures with the same parameters

**Table 4.** Comparison of predictive performance between HemolyticPredictor and existing hemolytic activity models

| Method | ACC | SEN | PRE | F1-score | MCC |
| --- | --- | --- | --- | --- | --- |
| SVM-Hem | 73.00% | 58.00% | 72.00% | 64.25% | 0.4400 |
| RF-Hem | 77.00% | 60.00% | 81.00% | 68.94% | 0.5300 |
| RNN-Hem | 76.00% | 76.00% | 70.00% | 72.88% | 0.5200 |
| AMPDeep | 79.97% | 83.28% | 79.88% | 81.54% | 0.5972 |
| HemoFinder | 81.67% | 81.67% | 81.79% | 81.73% | 0.6336 |
| **HemolyticPredictor** | **84.49%** | **82.67%** | **87.46%** | **85.00%** | **0.6909** |

#### Structural Analysis of Generated AMPs

To evaluate the structural characteristics of generated AMPs, we predicted their 3D structures using AlphaFold2 ^30^. Structural analysis revealed significant conformational diversity in the AMPs generated by our model, primarily comprising three stable motifs: canonical α-helical conformations, β-sheet topologies, and linear extended configurations.

Comparison of six representative generated AMPs (S1–S6) with two verified natural AMPs (R1, R2 from the APD3 database) highlighted key structural similarities (**Fig. 2A**). Structures S1–S3 adopt regular α-helices similar to R1, with helical parameters (e.g., pitch, turn angle) typical of AMPs, while S4–S6 exhibit extended conformations common in membrane-penetrating AMPs. To quantitatively assess prediction reliability, we analyzed AlphaFold2_’_s pLDDT scores. These scores (0–100) estimate confidence in predicted residue coordinates (higher scores denote greater reliability). generated AMPs had an average pLDDT of 66.47 ± 5.6. Notably, residues in key potential functional domains generally scored >70. These pLDDT scores support the reliability of the predicted structures and our model_’_s capacity to design AMPs with appropriate functional conformations. Furthermore, higher-confidence regions often corresponded to putative antibacterial domains, suggesting likely biological activity.

**Fig. 2:**
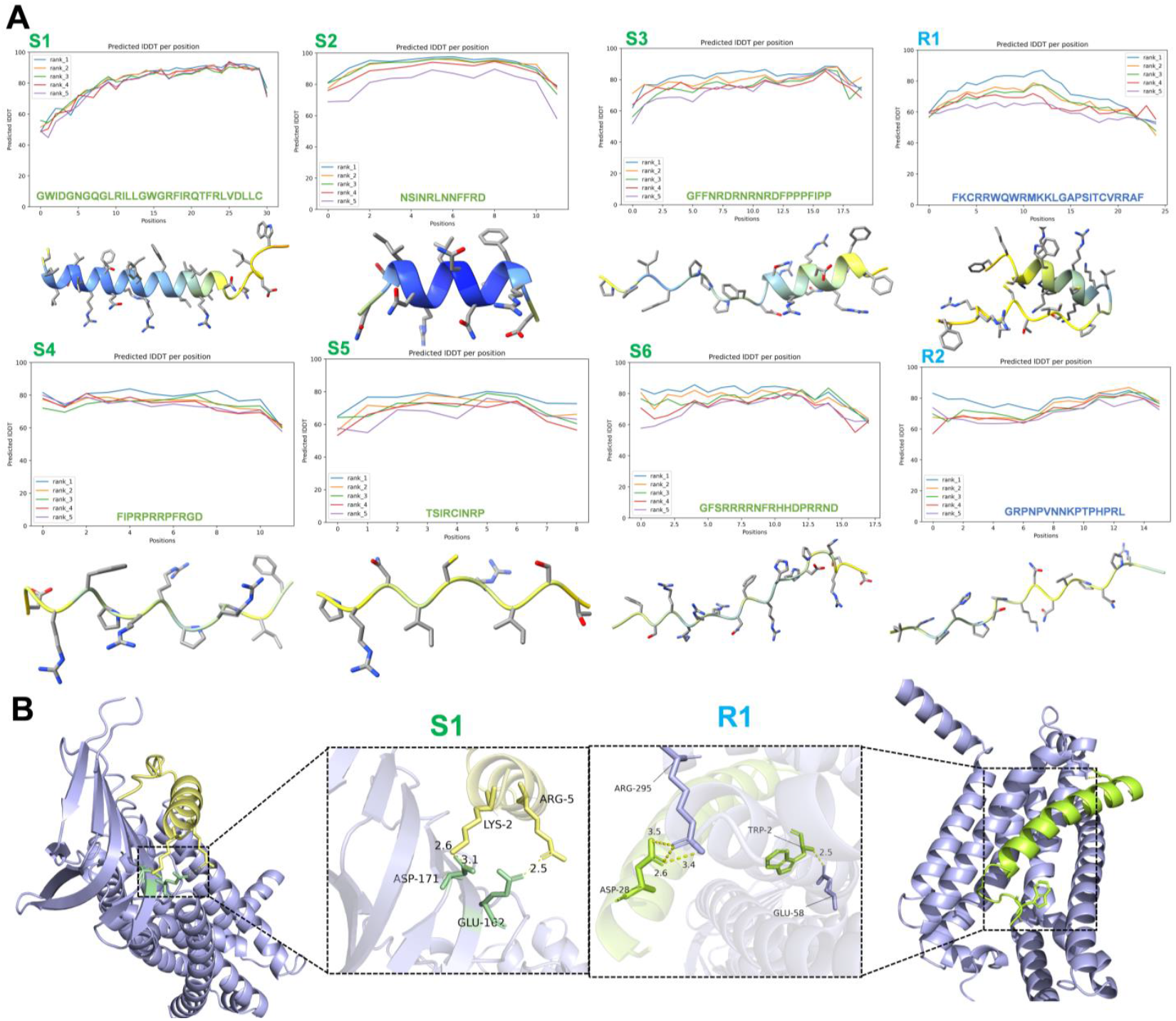
Analysis of generated AMP quality. (**A**) AlphaFold2-predicted structures of generated AMPs S1-S6 and reference AMPs R1, R2 from APD3, with pLDDT confidence plots. (**B**) Docking of peptide S1 with lipopolysaccharide (LPS): reference R1 (yellow), generated S1 (green), peptidoglycan (blue-purple); hydrogen bonds (yellow dashes), interacting residues (bold lines).

#### Peptide-Docking Analysis

To assess the therapeutic potential of generated peptides, we performed molecular docking simulations against lipopolysaccharide, a critical outer membrane component of Gram-negative bacteria ^31^, using the reference peptide R1. We quantified interaction energetics with ZDOCK ^32^,and visualized interfacial residue interactions using PyMOL to refine binding poses. Peptides generated by our model exhibited binding efficacy comparable to R1 **(example in Fig. 2B)**, supporting the potential of our generative approach. Docking results for an extended set of generated peptides, including comparative results for peptides from a suboptimal model, are provided in **Fig. S1**. These findings highlight the capability of our model to produce peptides with promising target engagement characteristics.

### Multi-Attribute Functional Analysis

To systematically evaluate the therapeutic potential of generated peptides, we employed our multi-attribute prediction framework (**Fig. S2**). The blue areas indicate that the peptide segments possess the corresponding biological activities. Notably, the peptides generated by the model not only exhibit structural features similar to those of natural AMPs, but also show highly consistent functional characteristics with real AMPs in key indicators such as the antibacterial spectrum (including inhibitory activities against Gram-positive and Gram-negative bacteria). These results suggest that our generation model can effectively produce structurally reasonable and functionally diverse antimicrobial peptides with physicochemical properties and biological activities comparable to those of natural AMPs.

### Screening Pipeline for Highly Active and Druggable AMPs Analysis

#### Efficacy Evaluation of an MIC Ranking Module and Regression Prediction Models

To assess the performance of our MIC ranking and regression models, we trained various machine learning algorithms— specifically, XGBoost, LightGBM, CatBoost, RandomForest, SVR, and Ridge—using datasets for four bacterial species: *S. aureus, C. albicans, E. coli*, and *P. aeruginosa*. Due to space constraints, we present detailed results for *S. aureus* here, with comprehensive comparisons for *C. albicans, E. coli*, and *P. aeruginosa* provided in **Figs. S3-S5**.

**Figs. 3A-B** demonstrates a positive correlation between the predicted MIC rankings and the actual labels for *S. aureus*, validating the effectiveness of our ranking models. Among these, LightGBM exhibited the best performance. In the screening pipeline, we utilize the top-performing ranking model to select the 500 most promising peptides. These selected peptides are then subjected to precise MIC prediction using our regression model, thereby optimizing the efficiency of subsequent wet-lab experiments.

**Fig. 3:**
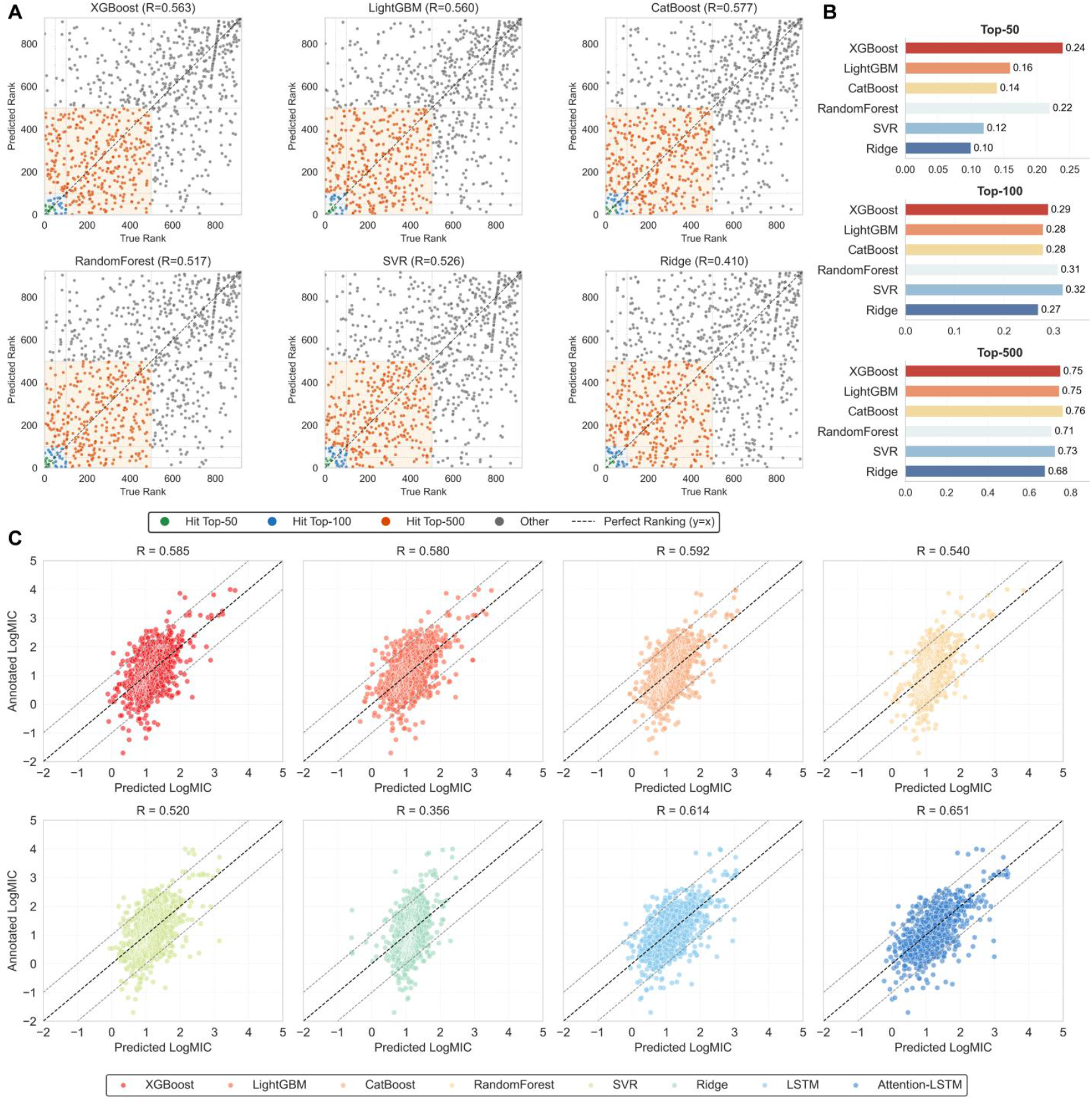
Comprehensive evaluation of machine learning models (including XGBoost, LightGBM, CatBoost, RandomForest, SVR, and Ridge) for ranking AMP activity against *S. aureus*. **(A)** A scatter plot assesses model performance in predicting AMP activity rankings across Top-K thresholds. Green dots signify correct identification of peptides within the actual top 50 active peptides, blue within the top 100, and orange within the top 500. This color-coded distribution enables a visual comparison of predictive efficacy across varying stringency levels. **(B)** A bar chart quantifies accuracy for Top-50, Top-100, and Top-500 predictions, where bar height reflects the proportion of peptides accurately ranked among the actual top K active peptides. Comparison of bar heights provides precise insight into model performance at different ranking depths. **(C)** Assessment of regression models for predicting MIC of AMPs against *S. aureus*. This scatter plot compares predicted MIC values to annotated ones for the models trained via regression. The solid line represents the ideal *y* = *x* correlation, while the dashed lines enclose predictions with absolute errors within the [-1, 1] range, indicating robust accuracy.

For the regression task, we trained models using Mean Squared Error loss on the training datasets for each bacterial species. We benchmarked our models against several established architectures, including the state-of-the-art LSTM-seq ^33^. As shown in **Fig. 3C** for *S. aureus* and in **Figs. S6-S8** for the other species, our Attention-LSTM model consistently outperformed all other models in terms of prediction accuracy and Pearson correlation coefficient.

#### Comparison of HemolyticPredictor Predictive Performance

Accurate prediction of hemolytic activity in AMPs is crucial for their development as safe therapeutics, as many AMPs can lyse mammalian erythrocytes, leading to hemoglobin release. To address this, we developed HemolyticPredictor, a model that leverages machine learning techniques from pattern recognition and bioinformatics to predict hemolytic potential. We benchmarked its performance against HemoFinder ^34^, the current state-of-the-art, and other established methods, using identical training and test datasets to ensure fairness. To ensure a fair comparison, all models were trained on the same training dataset and evaluated on the same independent test dataset ^35^. The detailed performance comparison results are presented in **Table 2**. HemolyticPredictor achieved superior performance with an accuracy of 84.49% and a MCC of 0.6909, demonstrating its effectiveness. This advancement aids in the design of AMPs with enhanced druggability by identifying candidates with minimal hemolytic activity.

#### Druggability Metric Screening of AMPs

We generated 10,000 novel peptide sequences de novo using our proposed PACD model. A comprehensive multi-attribute prediction initially determined the antimicrobial spectrum of each generated peptide, enabling the selection of candidates with putative activity against Gram-positive bacteria, Gram-negative bacteria, and fungi. Subsequently, a targeted hierarchical screening identified promising candidates for specific pathogens; for instance, potential *S. aureus*-targeting AMPs, predicted as active against Gram-positive bacteria, were ranked using an *S. aureus*-specific machine learning model. For each of the four key pathogens—*S. aureus, C. albicans, E. coli*, and *P. aeruginosa*—the top 500 sequences exhibiting the highest predicted activities were advanced for detailed druggability evaluation.

Druggability property distributions for these top 500 AMPs per pathogen are illustrated in **Figs. 4A-F**, approximately 70% of these sequences met predefined thresholds for individual druggability attributes. Applying a more stringent selection, sequences targeting *S. aureus* that simultaneously satisfied all druggability criteria were identified, resulting in 102 candidates (**Fig. 4G**). Analogous multi-criteria selections for the other pathogens (**Figs. S9–S11**) yielded sets of representative sequences (**Tables S2–S5**). This stringent filtering process inherently navigates the trade-offs among various druggability parameters, such as balancing low hemolytic activity with adequate stability and minimal cytotoxicity.

**Fig. 4:**
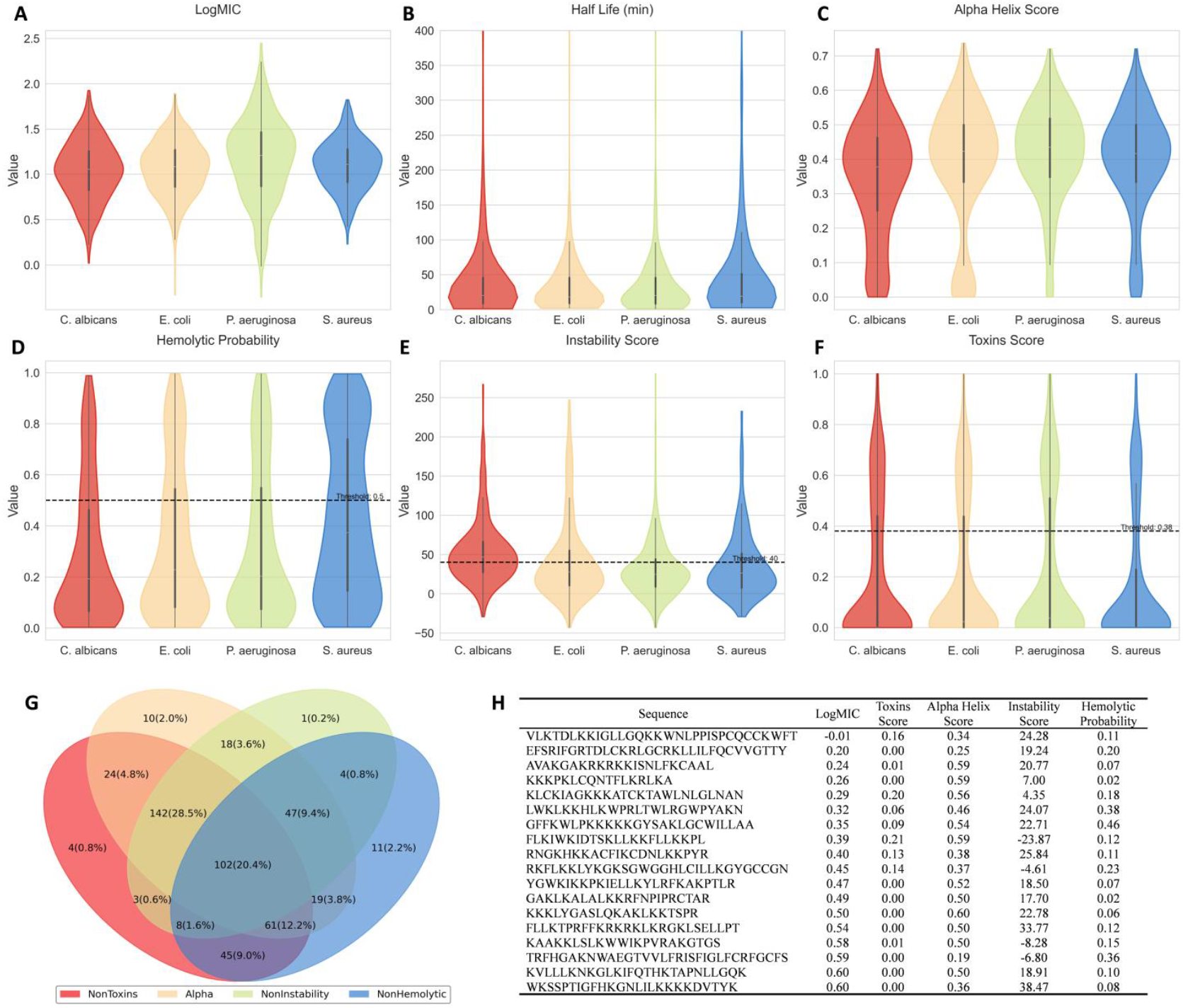
Distribution of predicted druggability properties for the AMPs. **(A)** Distribution of predicted logarithmic Minimum Inhibitory Concentration (LogMIC). **(B)** Distribution of predicted half-life (t_1/2_). **(C)** Distribution of predicted α-helical content. **(D)** Predicted hemolytic probability, where values below 0.5 signify non-hemolytic behavior and those ⩾ 0.5 suggest hemolysis. **(E)** Distribution of instability index, with scores ⩽40 denoting structural stability and scores>40 indicating instability. **(F)** Distribution of predicted toxicity scores, where values<0.38 imply non-toxicity, while values ⩾0.38 hint at potential toxicity. Together, these metrics offer a robust evaluation of AMP pharmacological properties profiles. **(G)** Venn diagram illustrating AMPs targeting *S. aureus* that satisfy the druggability screening criteria. **(H)** Detailing the drug-like properties of representative AMPs targeting *S. aureus*

Exemplary *S. aureus*-targeting AMPs that successfully passed this entire screening pipeline, thereby meeting all druggability standards, are presented in **Fig. 4H**. These selected candidates exhibit high predicted antimicrobial efficacy (MIC<10µM), negligible predicted toxicity, a propensity for α-helical secondary structure, low instability indices, and minimal hemolytic probabilities. These collective characteristics underscore the PACD model_’_s effectiveness in generating promising AMP candidates suitable for experimental validation and progression towards potential drug development.

#### Large-Scale Screening and Cross-Dataset Validation

To validate the efficacy of our screening pipeline, we applied it to an independent dataset of 43,000 human proteome peptide fragments. This dataset was previously analyzed by Torres et al.^36^, who identified 2,673 unique cAMPs using a scoring function and threshold-based filtering, with 20 of these subsequently confirmed for antimicrobial activity through experimental validation. Our pipeline first identified potential AMPs from these fragments, then predicted their antimicrobial spectra against Gram-positive bacteria, Gram-negative bacteria, and fungi. Subsequently, an optimized ranking model selected the top 500 peptides based on predicted activity against four key pathogens—*S. aureus, C. albicans, E. coli*, and *P. aeruginosa*. These selected peptides then underwent a thorough evaluation of their druggability properties, yielding the top 100 candidates for further analysis.

Analysis of the top 500 peptides (druggability property distributions are presented in **Fig. S12**) revealed that approximately 80% exhibited predicted MIC values below 10 µM. Most were also predicted to adopt α-helical secondary structures, exhibit high stability, and show low predicted toxicity. However, a significant proportion displayed potential hemolytic activity, which limited the number of candidates satisfying all druggability criteria. Specific sequences detailed in **Tables S6–S9** for each microbial species.

## Discussion

AMP-D3 constitutes a novel framework that integrates contrastive diffusion modeling with multimodal fusion to address critical challenges in AMP development. Specifically, it enables de novo peptide generation, precise recognition, detailed multi-attribute profiling, and systematic screening to pinpoint AMPs with elevated activity and druggability. Empirical assessments confirm AMP-D3_’_s exceptional performance in both identifying and generating AMPs. Additionally, the framework supports multi-task prediction across 22 biological activities and incorporates a tailored pipeline to evaluate the druggability of prospective AMPs, thereby expediting the identification of viable therapeutic candidates.

The framework substantially refines the design and selection of therapeutic AMPs through several innovative contributions, as follows. (1). CAST: This model enhances AMP recognition by merging deep embeddings with raw sequence data, significantly improving the detection of nuanced sequence patterns. (2). PACD: By incorporating biological priors, PACD increases the specificity and variety of generated AMPs, supporting their novelty and functional relevance. (3). Multi-attribute AMP prediction: This structure predicts 22 distinct biological properties, such as antifungal and anticancer effects, offering a comprehensive evaluation of each peptide_’_s capabilities. (4). A systematic screening pipeline for highly active and druggable AMPs: This systematic process assesses MIC, hemolysis, stability, and toxicity against targeted pathogens, facilitating the selection of AMPs with optimal activity and druggability. Together, these advancements position AMP-D3 as an effective tool for the targeted design and screening of AMPs, particularly in isolating candidates with improved druggability profiles, thus accelerating progress in AMP research. Nevertheless, two limitations require attention. Firstly, the framework underutilizes multimodal inputs, such as AMP structural data, during generation. This gap may compromise the folding stability of some peptides due to absent structural constraints. Secondly, the accuracy of multi-attribute prediction, a cornerstone of the screening process, remains open to refinement. Future work will prioritize integrating structural features into the generation phase and enhancing prediction precision across all attributes, thereby improving AMP-D3_’_s contribution to advancing AMP development.

## Methods

### PACD Model: Contrastive Diffusion AMP Generation

This section details the PACD model (**Fig. 5**), which integrates discrete diffusion modeling with contrastive learning to optimize AMP generation.

**Fig. 5:**
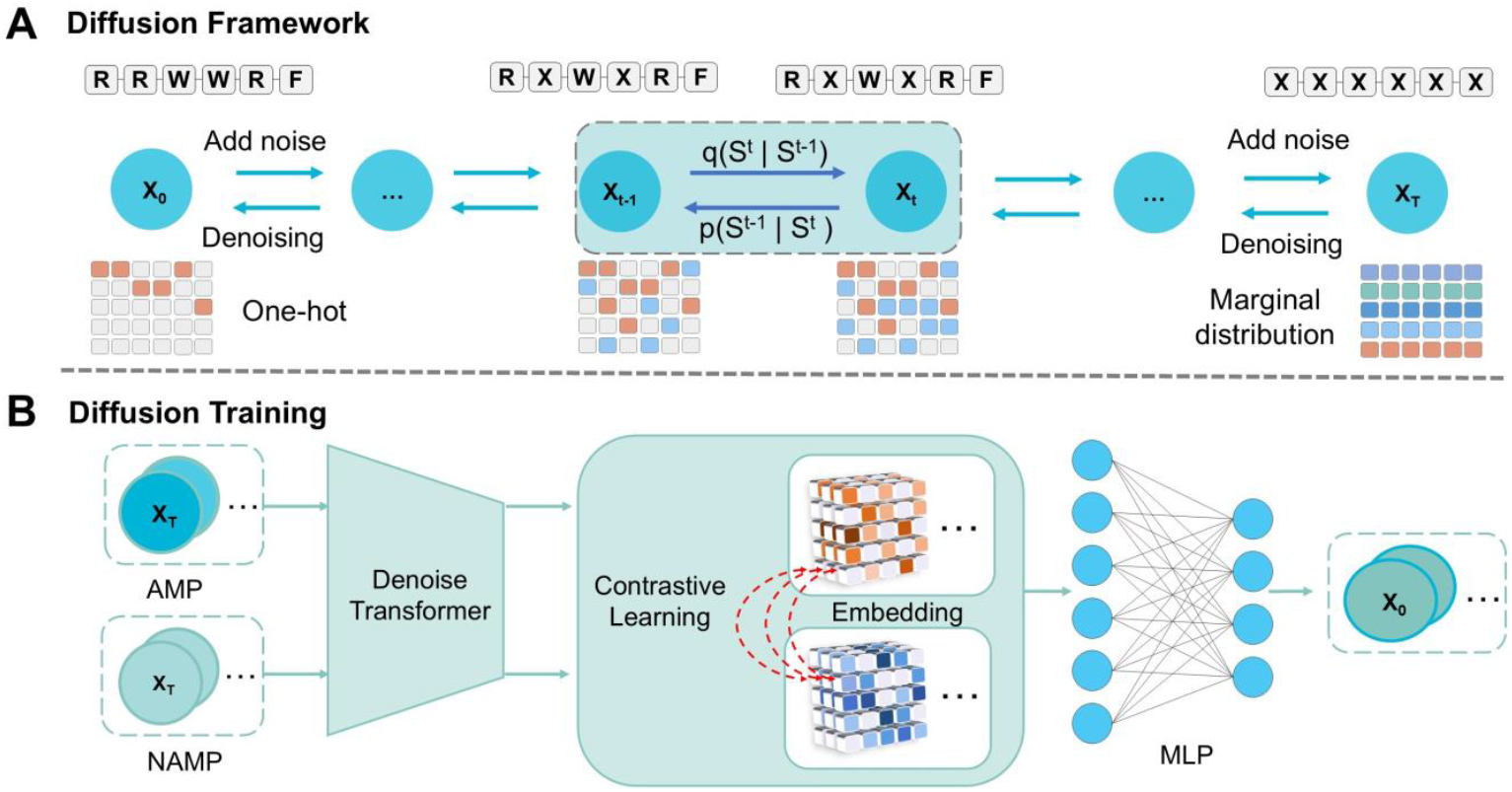
Overall workflow of the proposed PACD model. **(A)** Diffusion Framework: The diffusion process adopts a typical forward noise addition and reverse denoising mechanism, gradually perturbing the data distribution through a Markov chain and learning the reverse transformation using a parameterized network; **(B)** Diffusion Training: the Denoise Transformer extracts sequence features, combines a noise prediction head based on MLP to achieve precise denoising, and simultaneously introduces a contrastive learning objective to distinguish the latent representations of AMPs from non-antimicrobial peptides;

#### Diffusion Framework for AMP Generation

Our proposed PACD model **(Fig. 5A**) is an AMP generation framework. It significantly enhances the functional diversity of generated peptides by integrating discrete diffusion models for sequences ^37^ with contrastive learning ^38^. Inspired by Wang & Liu ^39^, PACD adapts diffusion probabilistic models ^37^ for peptide sequence generation.

The diffusion model achieves sequence generation through a defined Markov chain, comprising two key processes: forward diffusion and reverse generation ^40^. (1) Forward process: gradually injects noise into the authentic polypeptide sequence *X*_*0*_ = *S*_*0*_ to produce latent variable sequences *X*_*t*_ = *S*_*t*_. The noise injection is governed by transition probability *q*(*X*_*t*_ | *X*_*t-*1_) until the sequence completely degenerates into random noise distribution at timestep *T*. (2) Reverse generation process: reconstructs polypeptide sequences conforming to the real data distribution through gradual denoising via the learnable conditional probability (*X*_*t-*1_ | *X*_*t*_). This process, based on thermodynamic system evolution principles ^41^, ensures the structural rationality and functional potential of generated sequences.

Following Anand & Achim ^37^, we treat residue types as categorical data represented by one-hot encoding. During the forward process, we employ transition matrices to inject noise based on marginal distributions ^39^, progressively degrading sequences into random distributions. The reverse process is parameterized by conditional probability *q*(*S*_*t*-1_ | *S*_*t*_, *S*^*0*^) and implemented through a neural network that predicts the probability distribution of original sequence X_0_ ^42^. The key equations are given below:

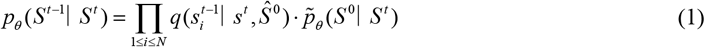

where *s*^*t*^ represents the one-hot feature of the *i*-th residue in sequence *S* at timestep *t*, and *S*^0^ denotes the model-predicted probability distribution of original sequence *S*^0^. We design the specific form of as:

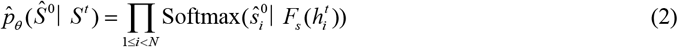

where 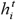 represents the input features of residue *i* with diffusion noise at timestep *t. F*_*s*_ is a hybrid neural network that first predicts residue-type noise from marginal distributions, then removes the noise to compute 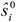 probabilities. Softmax is applied across all residue types. Our implementation *F*_*s*_ combines a Denoise Transformer with an MLP, the former learns contextual embeddings of residues from sequences, while the latter maps these embeddings to residue-type noise.

The embeddings of the generated sequences are subsequently refined through downstream contrastive learning. This integration (**Fig. 5B**) allows PACD to effectively capture sequence patterns related to therapeutic function, further enhancing generation accuracy and efficiency. Detailed computations are provided in **Supplementary Note 4**.

#### Loss Function

The diffusion process is optimized through dual objective components. First, sequence reconstruction fidelity is enforced via Kullback-Leibler divergence between predicted and ground-truth residue distributions at each timestep *t*:

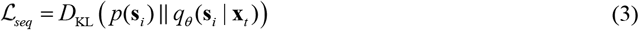

where *p*(**s**_*i*_) and *q*_*θ*_ (**s**_*i*_ | **x**_*t*_) represents the ground-truth probability and model-predicted distribution.

Then, the final composite objective function combines the sequence loss ℒ_*seq*_ and contrastive loss ℒ _*contrast*_ through weighted summation:

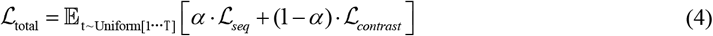

where *α* serves as a hyperparameter balancing task contributions, and Uniform[1, …, T] indicates uniform distribution over diffusion timesteps.

#### Denoise Transformer

this module processes two distinct input types: (1) data *X*_*t*_ generated through gradual Gaussian noise injection in the forward process of diffusion models, and (2) temporal embeddings for step *t*. Here, *X*_*t*_ is embedded using standard sinusoidal-cosine positional encoding ^43^, while timestep *t* generates conditional tokens through independent embedding.

To simultaneously enhance global feature extraction and preserve local amino acid characteristics, we employ Patchify Attention ^44^, a dual-mechanism architecture comprising global attention and local attention components.

Global Attention (sequence level): captures complex inter-residue relationships across protein sequences. For each head *i* in the h-head mechanism:

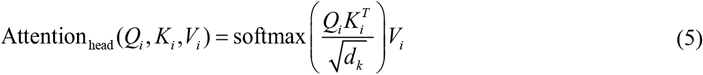

where 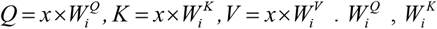, and 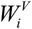 are head-specific weight matrices; *d* _*k*_ dimension of key vectors. The global attention output integrates multi-head representations through concatenation.

Local Attention (residue level): processes sequence segments of length *L* divided into patch-size patches. For each patch *j*:

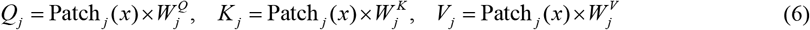

Attention weights follow the scaled dot-product calculation in Equation 6. Final local outputs are reconstructed through patch aggregation Local-Attention(*x*)=concat(*Patch*_*1*_, …, *Patch*_*p*_).

This architecture effectively captures hierarchical dependencies - global sequence patterns and local residue interactions - while enabling dynamic normalization parameter adjustment through conditional information integration, substantially enhancing model expressiveness.

### CAST Model: Cross-Attention AMPs Recognition

The proposed CAST architecture, depicted in **Fig. 6**, integrates evolutionary-scale protein modeling with multi-modal feature fusion for enhanced AMP identification.

**Fig. 6:**
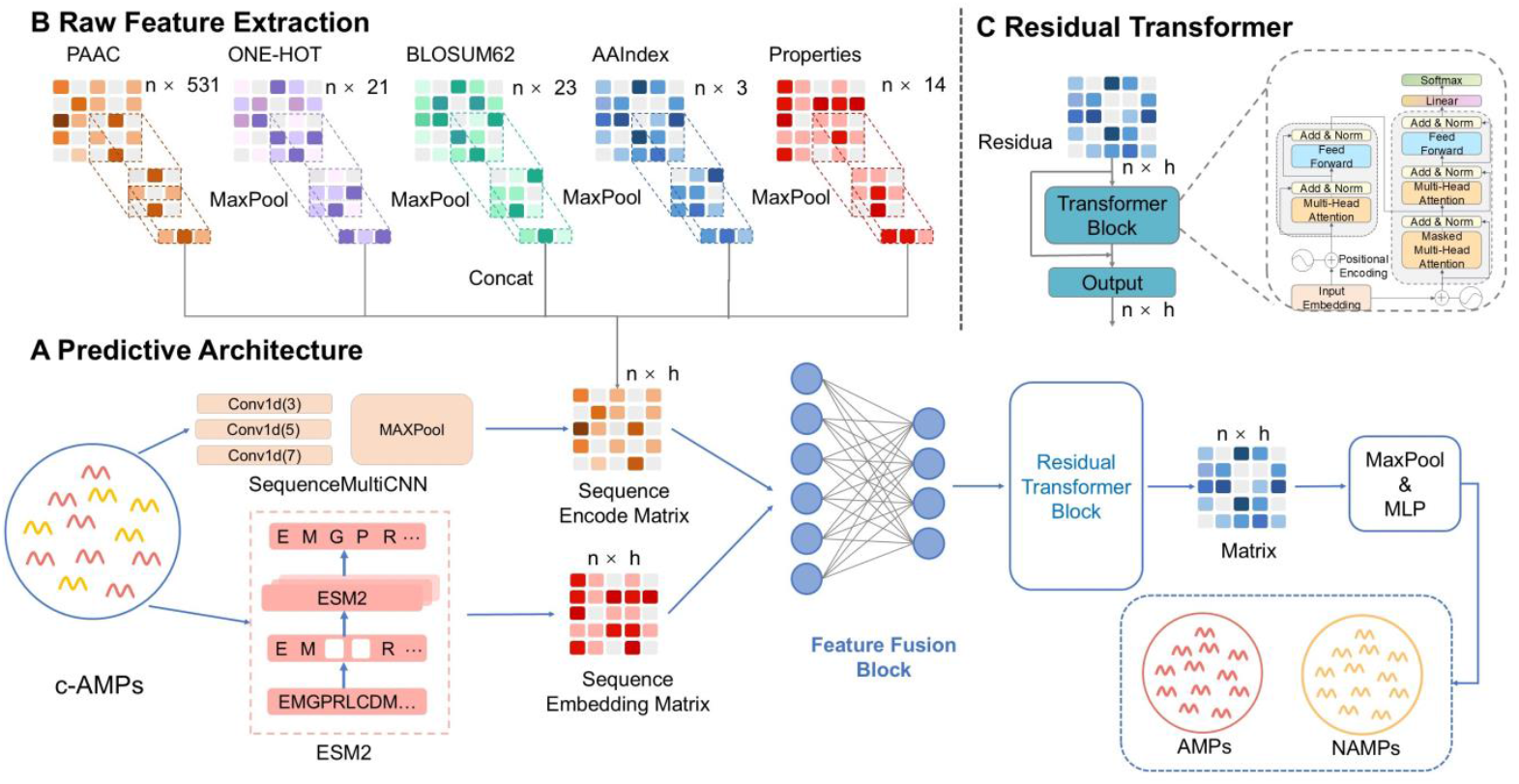
CAST recognition architecture. **(A)** Predictive Architecture: The input sequence undergoes dual feature extraction - high-dimensional embeddings are derived through the ESM2 model, while raw sequence features are extracted via SequenceMultiCNN. **(B)** Raw Feature Extraction: Integrates multi-dimensional raw features (including PAAC, One-hot, BLOSUM62, AAIndex, and physicochemical properties processed by MultiCNN into an encoded matrix) with semantic embeddings from ESM2. **(C)** Residual Transformer: A residual Transformer encoder with skip connections mitigates gradient vanishing, followed by MLP layers for final AMPs_’_ activity prediction.

#### Data Preprocessing

CAST transforms input sequences into dual complementary representations: High-Dimensional Embeddings: extracted using ESM2 ^19^, which learns universal protein representations through large-scale pretraining on billions of protein sequences. The esm2_t12_35M_UR50D model has demonstrated particular effectiveness for antimicrobial peptide prediction ^18^. Each peptide sequence is encoded as a 1 × 30 × 480 tensor:

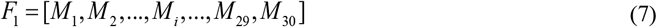

where *M*_*i*_ denotes the 480-dimensional embedding vector for the *i*-th residue.

Local Features: comprise five complementary amino acid representations (**Fig. 6B**): One-hot encoding *S*_1_ ; BLOSUM62 ^45^ matrix encoding *S*_2_; AAIndex ^46^ physicochemical profiles *S*_3_; Physicochemical property encoding *S*_4_; Pseudo-amino acid composition (PAAC) ^47^ *S*_5_. Please refer to **Supplementary Note 5** for details.

Employing raw feature representations enhances the breadth of input sequence matrix encoding. However, excessive redundant information may impair subsequent feature fusion and hinder Transformer performance. To address this limitation, inspired by multi-scale strategies ^15^, we implement a multi-kernel approach (3×3, 5×5, and 7×7 kernels) for each raw feature type to strengthen cross-scale feature extraction. The processing pipeline sequentially applies convolutional operations, batch normalization, and max pooling to improve computational efficiency and network robustness. The processed features are then concatenated to form comprehensive local sequence representations *F*_2_=Concat(*C*_1_, *C*_2_, *C*_3_, *C*_4_, *C*_5_), where *F*_2_ denotes the integrated local feature matrix.

#### CAST Architecture

As shown in **(Fig. 6A)**, CAST consists of four core modules: (1) Long-range dependency modeling module, which uses ESM-2 to extract long-range dependencies and semantic relationships in the sequence, thereby generating high-dimensional embedding features; (2) Multi-scale local feature extraction module, as illustrated in **(Fig. 6B)**, which extracts multi-scale sequence local features from the original features by adopting CNNs with different kernel sizes; (3) Cross-modal information fusion module, which integrates information from different modalities using a multi-attention mechanism; (4) Residual feature learning module, which includes a residual transformer block to enhance the feature capability.

#### Multi-Head Cross-Modal Dynamic Fusion

The fusion module synergistically integrates high-dimensional embeddings (*F*_1_) and local sequence features (*F*_2_) through a hybrid architecture combining multi-head attention with Feature-wise Linear Modulation ^48^. This dual-mechanism approach first projects input modalities into query (Q), key (K), and value (V) components via learnable linear transformations: *Q* _=_ *W*_*q*_ · *F*_1_, *K* _=_ *W*_*k*_ · *F*_2_, *V* _=_ *W*_*v*_ · *F*_2_, where *W*_*q*_, *W*_*k*_, *W*_*v*_ ∈ ℝ^input-dim×hidden-dim^ denote trainable parameter matrices. Subsequently, scaled dot-product attention computes interaction scores normalized by head dimension to stabilize gradients. The scaling factor head-dim stabilizes gradients, *F*_*fused*_ are fused feature matrix:

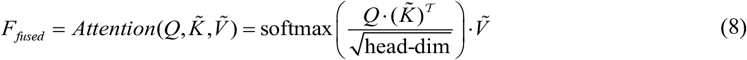

FiLM further enhances this process by dynamically modulating attention weights through feature-conditioned affine transformations. The resultant fused representation **(Equation 8)** effectively captures both global contextual patterns and localized biochemical constraints, particularly advantageous for AMP recognition requiring multi-scale feature integration.

### Multi-Attribute AMP Prediction

Our architecture employs parallel convolutional pathways (**Fig. 1C**) for simultaneous prediction of 22 biological activities. The pipeline initiates through four complementary encoding schemes: (1) one-hot sequence vectors, (2) BLOSUM62 substitution matrices, (3) AAIndex physicochemical profiles, and (4) PAAC. Each encoding undergoes independent processing via dedicated CNN branches equipped with multi-scale convolutional kernels (sizes 2-5), followed by max-pooling operations for spatial reduction. Following feature extraction, branch outputs are concatenated into a unified representation before final projection through three fully connected layers. The final prediction layer generates probability scores across therapeutic categories including antibacterial, antiviral, and anticancer activities. This architecture builds upon established methodologies demonstrating efficacy in multifunctional AMP characterization ^15^.

### A Systematic Screening Pipeline for Highly Active and Druggable AMPs

Despite feasible de novo generation and broad-spectrum screening of AMPs, comprehensively evaluating their pharmacological properties remains challenging. Our multi-stage screening pipeline (**Fig. 1E**) addresses this by efficiently identifying highly active AMPs against specific pathogens (e.g., *C. albicans, E. coli, P. aeruginosa*, and *S. aureus*), integrating activity prediction, ranking, and druggability assessment to refine candidate selection.

The pipeline, starting with 10,000 peptides from PACD, first uses broad-spectrum activity prediction (against Gram-positive/-negative bacteria and fungi) to reduce the initial set. A dedicated pre-screening module then ranks and selects the top 500 AMPs based on predicted activity against key bacteria. These candidates undergo rigorous multi-dimensional evaluation, focusing on:

- Low hemolytic activity: low hemolytic potential (predicted by HemolyticPredictor).
- High stability: instability Index < 40.
- Low toxicity: predicted by ToxinPred3.0.
- α-helical structure: predicted α-helical conformation (crucial for membrane disruption).

Peptides meeting these criteria are ranked by predicted MIC values from our regression model. The top 100 peptides (lowest predicted MICs, highest anticipated potency) further analysis based on predicted properties. This step further assesses their antimicrobial efficacy, safety (hemolysis, cytotoxicity), and stability, thereby supporting computational predictions and contributing to novel AMP discovery.

#### Antimicrobial Activity Ranking Pre-screening Module

This module leverages a Learning-to-Rank algorithm, trained on comparative MIC data, to predict peptide activity with precision. By emphasizing relative potency, it mitigates biases and experimental variability, ensuring robust performance. LightGBM underpins this step, chosen for its exceptional accuracy and adaptability.

#### Druggability Metric Screening Module

For AMPs to be viable clinical candidates, they must demonstrate not only robust antimicrobial efficacy but also essential pharmaceutical attributes ensuring safety and practicality. These attributes encompass stability, minimal hemolytic activity, and low cytotoxicity. To assess these properties comprehensively, we implemented a suite of computational models, culminating in a multi-faceted screening protocol.

#### Prediction of Antimicrobial Activity (MIC)

Inspired by Huang et al._’_s ^**33**^ AMP-MIC prediction framework, we developed an advanced regression model to predict the MIC of AMPs, building on prior LSTM-based frameworks. By integrating ESM2 protein language model embeddings to capture complex sequence features and an attention mechanism to emphasize critical regions for antimicrobial activity, our Attention-LSTM model achieves superior prediction accuracy and Pearson correlation compared to state-of-the-art methods.

#### Prediction of Hemolytic Activity

To predict hemolytic activity, a key safety parameter, we developed HemolyticPredictor (**Fig. S13**), a machine learning model that integrates five peptide descriptors: AAC, DPC, PAAC, CKSAAGP, and PHYC. These descriptors encode sequence information across various dimensions, such as composition, fragment patterns, and physicochemical properties, providing a robust feature set for classification. Utilizing the LightGBM algorithm, HemolyticPredictor accurately identifies peptides with potential hemolytic risks.

#### Assessment of Peptide Stability

Peptide stability significantly impacts pharmacokinetics and efficacy. We employed two metrics: (1) a regression model by Cavaco et al. ^49^ predicting half-life (*t*_1/2_, in minutes):

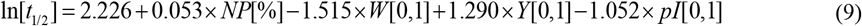

where NP is the percentage of nonpolar residues, indicates W presence (1 if present, 0 otherwise), Y indicates Tyr status (0 if absent or one, 1 if two or more), and pI is 0 if <10, 1 otherwise. (2) The Instability Index from Biopython, assessing sequence-based instability. Together, these provide a comprehensive stability evaluation.

#### Assessment of Peptide Toxicity

Peptide toxicity was evaluated using ToxinPred3.0 ^50^, a machine learning tool trained on diverse toxic/non-toxic peptides predicting toxicity based on sequence features.

## Acknowledgements

This work was supported by the National Natural Science Foundation of China (62372234, 62072243), the Natural Science Foundation of Jiangsu (BK20201304), Major and Seed Inter-Disciplinary Research project awarded by Monash University, and the Natural Science Research Start-up Foundation of Recruiting Talents of Nanjing University of Posts and Telecommunications (Grant No. NY223062), and the Natural Science Foundation of Nanjing University of Posts and Telecommunications (Grant No. NY224158).

## Ethics declarations

The authors declare that they have no competing interests.

## Supplementary Information

This PDF file includes:

Supplementary Notes 1 to 5

Supplementary Figs. S1 to S13

Supplementary Tables S1 to S9

